# Loss of AKAP1 triggers Drp1 dephosphorylation-mediated mitochondrial fission and loss in retinal ganglion cells

**DOI:** 10.1101/790139

**Authors:** Genea Edwards, Guy A. Perkins, Keun-Young Kim, YeEun Kong, Yonghoon Lee, Soo-Ho Choi, Yujia Liu, Dorota Skowronska-Krawczyk, Robert N. Weinreb, Linda Zangwill, Stefan Strack, Won-Kyu Ju

## Abstract

Impairment of mitochondrial structure and function is strongly linked to glaucoma pathogenesis. Despite the widely appreciated disease relevance of mitochondrial dysfunction and loss, the molecular mechanisms underlying mitochondrial fragmentation and metabolic stress in glaucoma are poorly understood. We demonstrate here that glaucomatous retinal ganglion cells (RGCs) show loss of A-kinase anchoring protein 1 (AKAP1), activation of calcineurin (CaN) and reduction of dynamin-related protein 1 (Drp1) phosphorylation at serine 637 (Ser637). These findings suggest that AKAP1-mediated phosphorylation of Drp1 at Ser637 has a critical role in RGC survival in glaucomatous neurodegeneration. Male mice lacking AKAP1 show increases of CaN and total Drp1 level, as well as a decrease of Drp1 phosphorylation at Ser637 in the retina. Ultrastructural analysis of mitochondria shows that loss of AKAP1 triggers mitochondrial fragmentation and loss, as well as mitophagosome formation in RGCs. Loss of AKAP1 deregulates oxidative phosphorylation (OXPHOS) complexes (Cxs) by increasing CxII and decreasing CxIII-V, leading to metabolic and oxidative stress. Also, loss of AKAP1 decreases Akt phosphorylation at Serine 473 (Ser473) and threonine 308 (Thr308) and activates the Bim/Bax signaling pathway in the retina. These results suggest that loss of AKAP1 has a critical role in RGC dysfunction by decreasing Drp1 phosphorylation at Ser637, deregulating OXPHOS, decreasing Akt phosphorylation at Ser473 and Thr308, and activating the Bim/Bax pathway in glaucomatous neurodegeneration. Thus, we propose that overexpression of AKAP1 or modulation of Drp1 phosphorylation at Ser637 are potential therapeutic strategies for neuroprotective intervention in glaucoma and other mitochondria-related optic neuropathies.

## Introduction

Primary open-angle glaucoma (POAG) is characterized by a slow and progressive degeneration of retinal ganglion cells (RGCs) and their axons, leading to loss of visual function^1^. The factors contributing to degeneration of the RGC and its axon degeneration in POAG are not well understood. Recent studies have shown that POAG patients have mitochondrial abnormalities^2–7^. Evidence from our group and others indicates that compromised mitochondrial dynamics, metabolic stress and mitochondrial dysfunction by glaucomatous insults such as elevated intraocular pressure (IOP) and oxidative stress are critical to RGC loss in experimental glaucoma^8–14^. Despite the widely appreciated disease relevance of mitochondrial dysfunction and loss, the molecular mechanisms underlying the impairment of mitochondrial structure and function in glaucoma are poorly understood.

A-kinase anchoring protein 1 (AKAP1), a scaffold protein, is part of the elaborate architecture which binds to the regulatory subunit of protein kinase A (PKA) along with other signaling molecules such as Src kinase, protein phosphatase 1 (PP1), Ca^2+^-dependent phosphatase calcineurin (CaN) and phosphodiesterase 4 (PDE4)^15–20^. By recruiting the PKA holoenzyme to the outer mitochondrial membrane (OMM), AKAP1 is strategically positioned to integrate multiple signal transduction cascades, including cyclic adenosine 3’5’-monophosphate (cAMP) signaling, in the regulation of mitochondrial shape and function. AKAP1 facilitates dynamin-related protein 1 (Drp1) phosphorylation and subsequent mitochondrial elongation. It also increases adenosine triphosphate (ATP) generation and mitochondrial membrane potential^21, 22^, and these collective changes are neuroprotective ^23, 24^. Emerging evidence indicates that AKAP1 protects from cerebral ischemic stroke by maintaining respiratory chain activity, inhibiting Drp1 dependent mitochondrial fission and superoxide production, and delaying Ca^2+^ deregulation ^24^.

Excessive mitochondrial fission-mediated dysfunction has been implicated in various neurodegenerative diseases including glaucoma^8, 11, 25, 26^. Complexes of Drp1 assemble from the cytosol onto mitochondria at focal sites of mitochondrial fission^27, 28^. Recent evidence indicates that post-translational modifications of Drp1 are linked to mitochondrial dysfunction-mediated bioenergetic failure, synaptic injury and neuronal cell death^25, 29–32^. Phosphorylation of Drp1 at serine 637 (Ser637) by the cAMP-dependent PKA promotes mitochondrial fusion, whereas dephosphorylation by the CaN promotes mitochondrial fission by decreasing Drp1 activity^33–35^. Previous studies showed that inhibition of Drp1 activity prevents mitochondrial fission and protects RGCs and their axons in experimental glaucoma^11, 36, 37^. However, it remains unknown whether AKAP1-mediated Drp1 phosphorylation at Ser637 play a critical role in glaucomatous neurodegeneration. Along this line, the existence of AKAP1 in RGCs and its role in glaucoma are completely unknown.

To address these questions, we investigated AKAP1 protein expression and Drp1 phosphorylation at Ser637 in retinas from glaucomatous DBA/2J mice, and also evaluated the effect of AKAP1 loss in the retina using AKAP1 knockout (AKAP1^-/-^) mice.

## Results

### AKAP1 deficiency in glaucomatous RGCs

We examined AKAP1 protein expression in the retina of a mouse model of glaucoma DBA/2J mice, which spontaneously develop elevated IOP and glaucomatous damage with age^8, 11, 38^.

Control DBA/2J-*Gpnmb^+^/SjJ* (D2-*Gpnmb^+^*) was used in which the gene for Gpnmb that has been altered to wild type sequence by targeted gene mutation. It is otherwise identical to the DBA/2J mouse strain, but maintains normal IOP during aging and does not develop optic nerve axon loss^39, 40^. Our results demonstrated for the first time that elevated IOP significantly decreased AKAP1 protein expression in the retinas of 10-month-old glaucomatous DBA/2J mice compared with the retina of age-matched control D2-*Gpnmb^+^* mice (Fig. 1a). We observed that AKAP1 immunoreactivity was highly present in the outer plexiform layer (OPL) and ganglion cell layer (GCL) in D2-*Gpnmb^+^* retina (Fig. 1b). More specifically, AKAP1 immunoreactivity was colocalized with neuronal class III β-tubulin (TUJ1)-positive RGCs in the GCL of D2-*Gpnmb^+^* retina. Of interest, however, AKAP1 immunoreactivity was decreased in the OPL and TUJ1-positive RGCs in the GCL of glaucomatous DBA/2J retina (Fig. 1b and c).

**Fig. 1.**
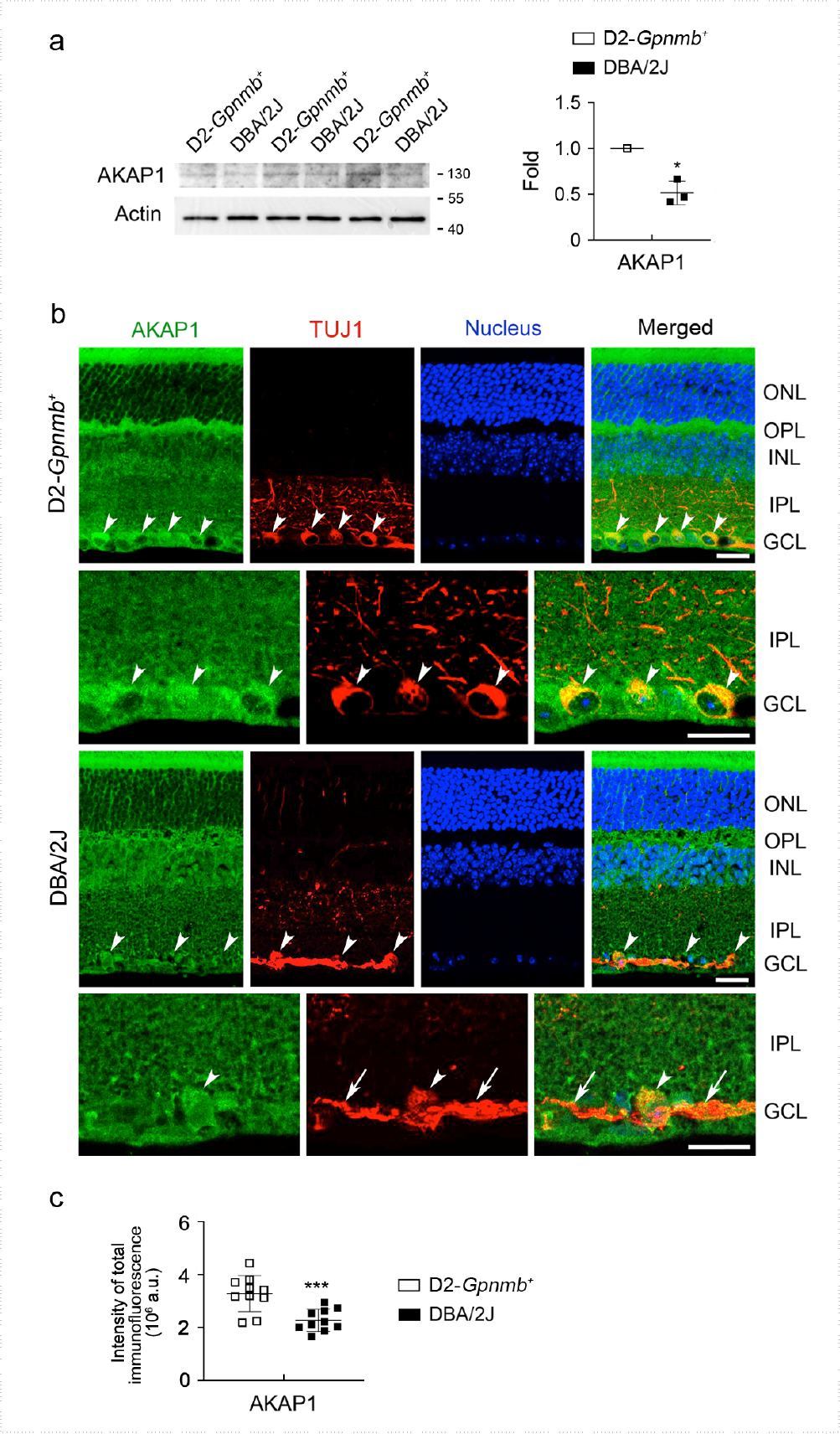
AKAP1 deficiency in glaucomatous RGCs. (a) Western blot analysis for AKAP1 in the retinas of 10-month-old glaucomatous DBA/2J and age-matched D2-*Gpnmb^+^* mice. (b) Representative images from immunohistochemical analyses for AKAP1 (green) and TUJ1 (red) in the retina of D2-*Gpnmb^+^* and glaucomatous DBA/2J mice. Arrowheads indicate accumulation of AKAP1 co-labeled with TUJ1 in RGC somas and arrows indicate TUJ1-labeled axon bundles. Note that glaucomatous RGCs showed decrease of AKAP1 protein expression. Blue color indicates nucleus. (c) Quantitative analysis for fluorescent intensity showed a significant decrease of AKAP1 immunoreactivity in the retina of glaucomatous DBA/2J mice. GCL, ganglion cell layer; IPL, inner plexiform layer; INL, inner nuclear layer; OPL, outer plexiform layer; ONL, outer nuclear layer. Mean ± SD; *n* = 3 (a) and *n* = 10 (c); *\*P* < 0.05 and *\*\*\*P* < 0.001 (two-tailed unpaired Student’s *t*-test). Scale bar: 20 µm.

### Activation of CaN and dephosphorylation of Drp1 at Ser637 in glaucomatous retina

AKAP1 binds with two Serine/Threonine phosphatases, PP1 and CaN^41, 42^. Loss of AKAP1 causes Drp1-mediated mitochondrial fission and decreases Drp1phsophorylation at Ser637 in neuronal cells of the brain^21, 23, 24, 43, 44^. More importantly, AKAP1 protects brain neuronal cells against cerebral ischemic stroke by inhibiting Drp1-dependent mitochondrial fission^24^. Since elevated IOP increased CaN and total Drp1 protein expression^11, 45^, as well as Drp1 inhibition rescued RGCs and their axons by preserving mitochondrial integrity in the retina and/or glial lamina of glaucomatous DBA/2J mice^11^, we examined the expression levels of CaN and total Drp1, as well as phosphorylation of Drp1 at Ser637 in the retina of 10-month-old glaucomatous DBA/2J mice. We observed a significant increase of CaN protein expression in glaucomatous DBA/2J retina (Fig. 2a). Consistently, our results showed an increase of CaN immunoreactivity in RNA-binding protein with multiple splicing (RBPMS)-positive RGCs as well as neurons in the inner nuclear layer (INL) of glaucomatous DBA/2J retina (Fig. 2b and c). We also observed a significant increase of total Drp1 protein expression but a significant dephosphorylation of Drp1 Ser637 in glaucomatous DBA/2J retina (Fig. 2d). Consistently, our results showed an increase of Drp1 immunoreactivity in TUJ1-positive RGCs of glaucomatous DBA/2J retina (Fig. 2e and f). These results suggest that elevated IOP-induced CaN activation is associated with dephosphorylation of Drp1 at Ser637 in glaucomatous RGCs, leading to mitochondrial fisison^11^.

**Fig. 2.**
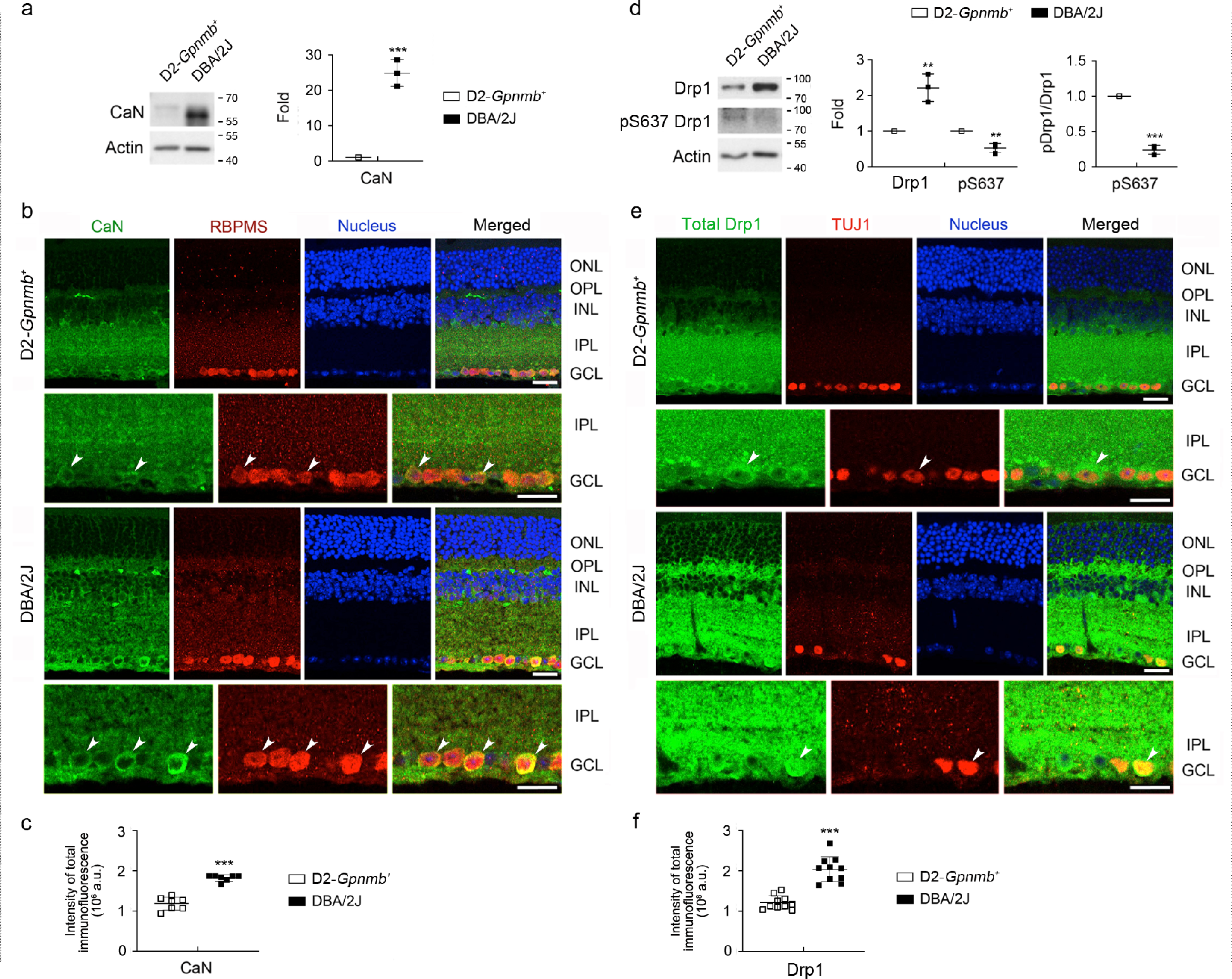
CaN-mediated dephosphorylation of Drp1 at S637 in glaucomatous retina. (a) Western blot analyses for CaN in the retinas of 10-month-old glaucomatous DBA/2J and age-matched D2-*Gpnmb^+^* mice. (b) Representative images from immunohistochemical analyses for CaN (green, arrowheads) co-labeled with RBPMS (red, arrowheads) in RGCs. Note that glaucomatous RGCs showed increases of CaN protein expression. Blue color indicates nucleus. (c) Quantitative analysis for fluorescent intensity showed a significant increase of CaN immunoreactivity in the retina of glaucomatous DBA/2J mice. (d) Western blot analyses for total Drp1 and phospho-Drp1 Ser637 in the retinas of glaucomatous DBA/2J and age-matched D2-*Gpnmb^+^* mice. (e) Representative images from immunohistochemical analyses for total Drp1 (green, arrowheads) co-labeled with TUJ1 (red, arrowheads) in RGCs. Note that glaucomatous RGCs showed increases of total Drp1 protein expression. Blue color indicates nucleus. (f) Quantitative analysis for fluorescent intensity showed a significant increase of Drp1 immunoreactivity in the retina of glaucomatous DBA/2J mice. GCL, ganglion cell layer; IPL, inner plexiform layer; INL, inner nuclear layer; OPL, outer plexiform layer; ONL, outer nuclear layer. Mean ± SD; *n* = 3 (a and d) and *n* = 10 (c and f); *\*\*P* < 0.01 and *\*\*\*P* < 0.001 (two-tailed unpaired Student’s *t*-test). Scale bar: 20 µm.

### Loss of AKAP1 triggers activation of CaN and dephosphorylation of Drp1 at Ser637 in the retina

We next used the retinas from mice lacking 10-month-old AKAP1 (AKAP1^-/-^)^24^ and age-matched wild-type (WT) mice and examined the expression level of CaN. Following the confirmation of a significant decrease of *AKAP1* gene expression (Fig. 3a), we observed a significant increase of CaN protein expression in AKAP1^-/-^ retina compared with WT retina (Fig. 3b). In addition, increased CaN immunoreactivity was present in RBPMS-positive RGCs as well as neurons in the INL in AKAP1^-/-^ retina (Fig. 3c and d). We further examined expression levels of AMP-activated protein kinase (AMPK) and Drp1 protein, as well as phosphorylation of AMPK at threonine 172 (Thr172) and Drp1 at Ser637 in AKAP1^-/-^ retina. There was no significant difference in the phosphorylation of AMPK at Thr172 between WT and AKAP1^-/-^ retinas, whereas we observed a significant decrease of total AMPK protein expression in AKAP1^-/-^ retina (Fig. 4a). Interestingly, we observed a significant increase of total Drp1 protein expression in in AKAP1^-/-^ retina (Fig. 4b). Also, AKAP1^-/-^ retina showed a significant dephosphorylation of Drp1at Ser637, whereas there was no significant difference in the phosphorylation of Drp1 at Ser616 between WT and AKAP1^-/-^ retinas (Fig. 4b). Consistent with this finding, increased Drp1 immunoreactivity was present in RBPMS-positive RGCs of AKAP1^-/-^ retina (Fig. 4c and d).

**Fig. 3.**
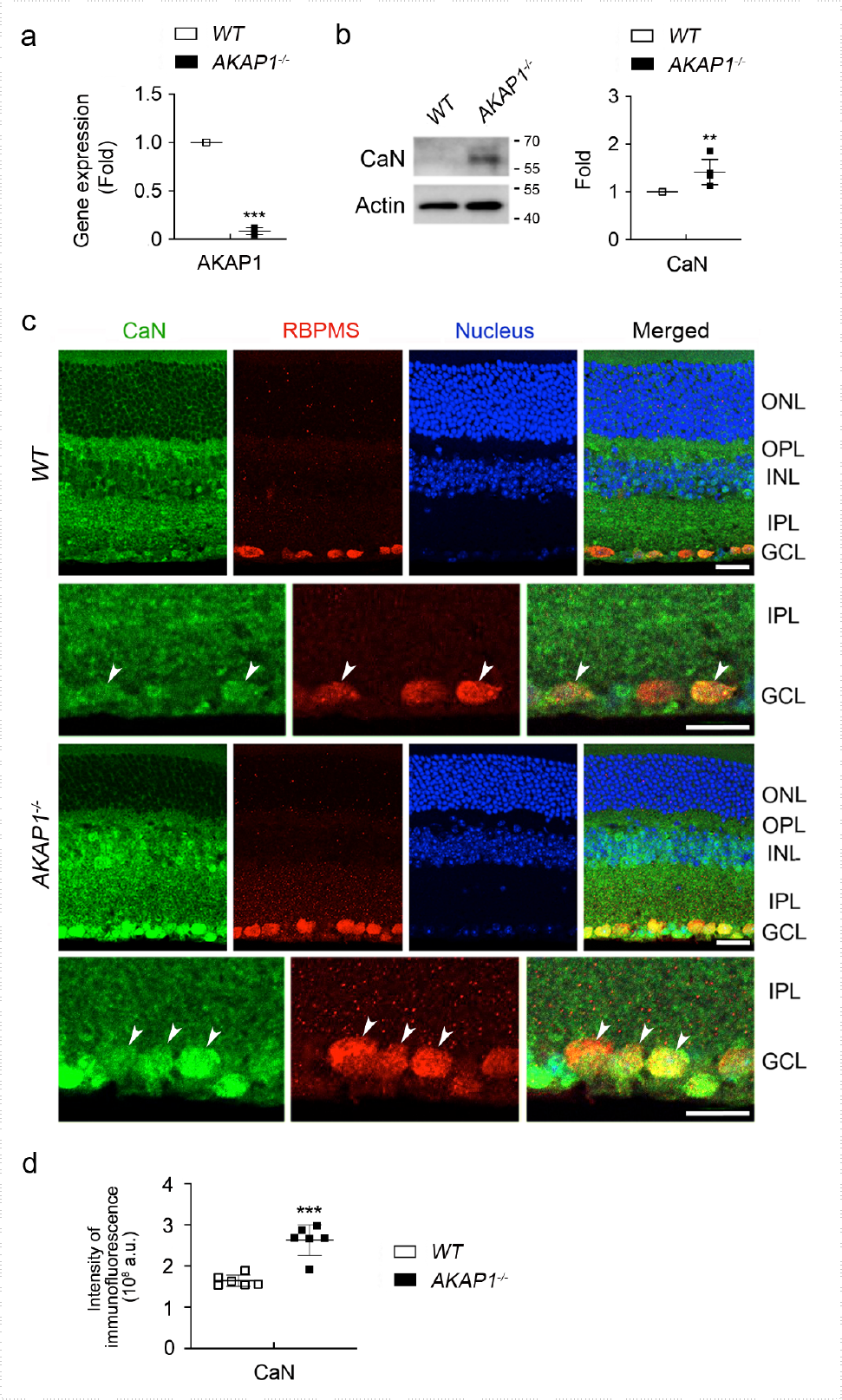
CaN expression in AKAP1^-/-^ retina. (a) Quantitative RT-PCR analysis for AKAP1 in the retinas of AKAP1^-/-^ and WT mice. (b) Western blot analysis for CaN in the retinas of AKAP1^-/-^ and WT mice. (c) Representative images from immunohistochemical analyses for CaN (green, arrowheads) co-labeled with RBPMS (red, arrowheads) in RGCs. Note that glaucomatous RGCs showed increases of CaN protein expression in the GCL and INL. Blue color indicates nucleus. (d) Quantitative analysis for fluorescent intensity showed a significant increase of CaN immunoreactivity in the retina of AKAP1^-/-^ mice. GCL, ganglion cell layer; IPL, inner plexiform layer; INL, inner nuclear layer; OPL, outer plexiform layer; ONL, outer nuclear layer. Mean ± SD; *n* = 3 or 4 (a and b) and *n* = 10 (d); *\*\*P* < 0.01 and *\*\*\*P* < 0.001 (two-tailed unpaired Student’s *t*-test). Scale bar: 20 µm.

**Fig. 4.**
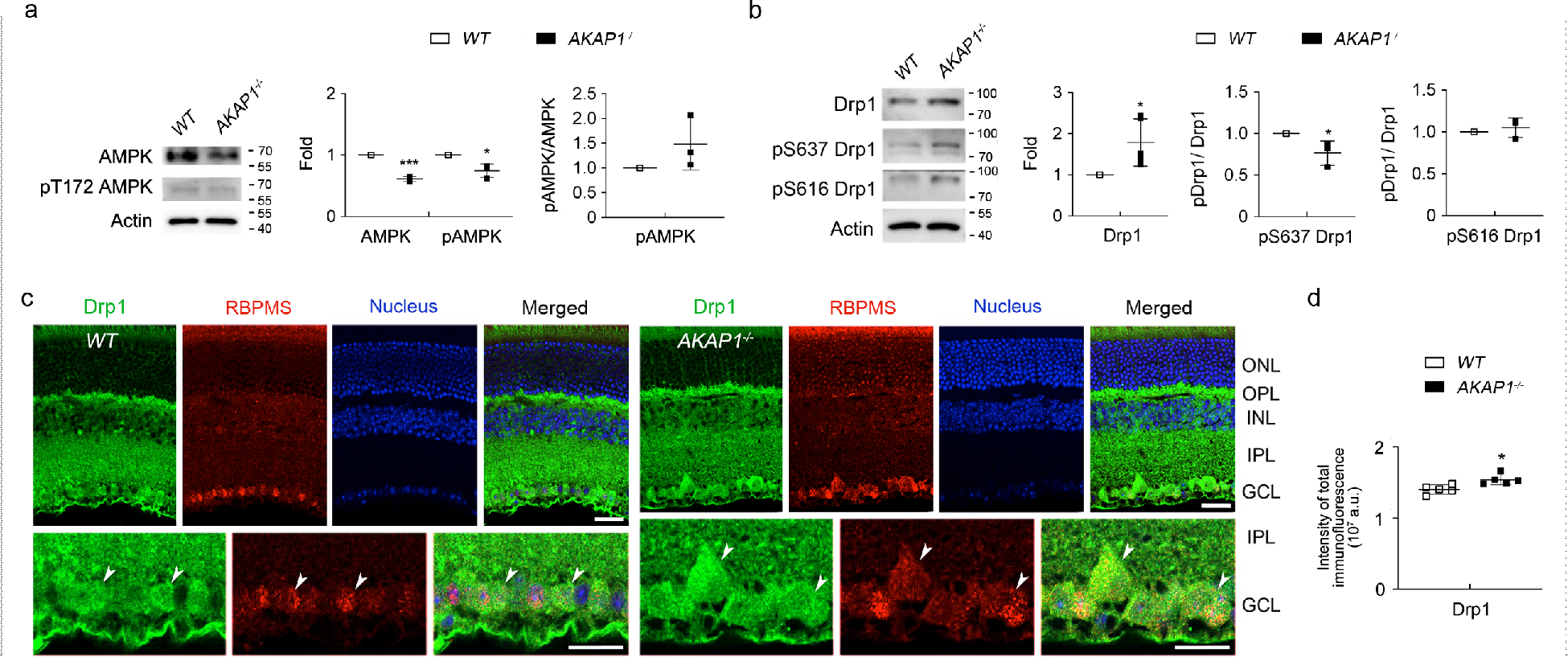
Phosphorylation of AMPK at Thr172 and Drp1 at Ser637 in AKAP1^-/-^ retina. (a) Western blot analysis for AMPK and phospho-AMPK Thr172 in the retinas of AKAP1^-/-^ and WT mice. (b) Western blot analysis for total Drp1, phospho-Drp1 Ser637 and phospho-Drp1 Ser616 in the retinas of AKAP1^-/-^ and WT mice. (c) Representative images from immunohistochemical analyses for total Drp1 (green, arrowheads) co-labeled with RBPMS (red, arrowheads) in RGCs. Note that glaucomatous RGCs showed an increase of total Drp1 protein expression in RGCs. Blue color indicates nucleus. (d) Quantitative analysis for fluorescent intensity showed a significant increase of Drp1 immunoreactivity in the retina of AKAP1^-/-^ mice. GCL, ganglion cell layer; IPL, inner plexiform layer; INL, inner nuclear layer; OPL, outer plexiform layer; ONL, outer nuclear layer. Mean ± SD; *n* = 3 or 6 (a) and *n* = 10 (d); *\*P* < 0.05 (two-tailed unpaired Student’s *t*-test). Scale bar: 20 µm.

### AKAP1 loss causes mitochondrial fission and mitophagosome formation in RGCs

We further examined the expression levels of optic atrophy type 1 (OPA1) and mitofusin (Mfn) 1 and 2 in AKAP1^-/-^ retina and the alteration of mitochondrial ultrastructure in AKAP1^-/-^ RGC somas. We observed no significant difference in *OPA1* gene expression, but a significant decrease of total OPA1 protein expression in AKAP1^-/-^ retina compared with WT (Fig. 5a).

**Fig. 5.**
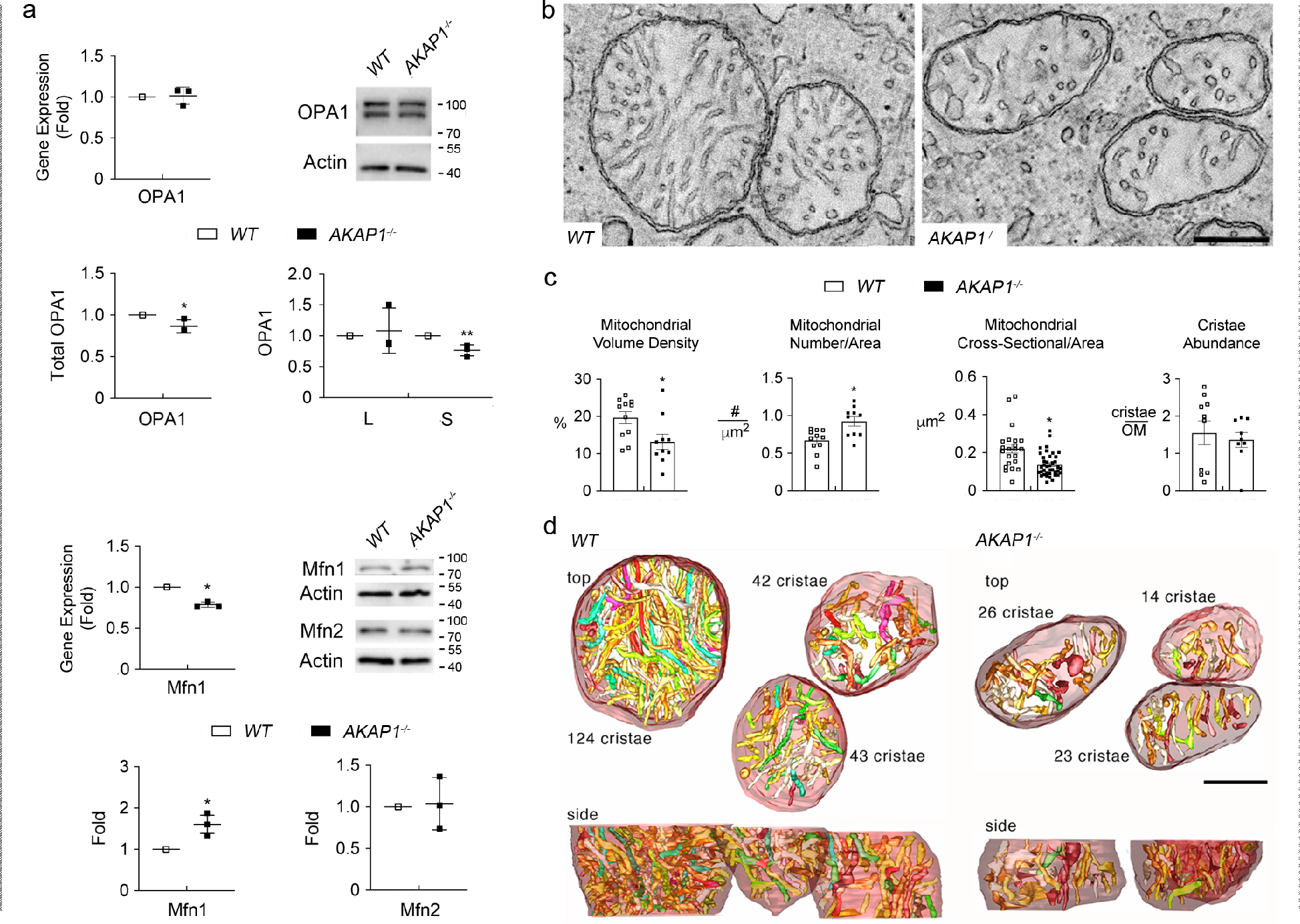
Mitochondrial fission in AKAP1^-/-^ RGC somas. (a) Quantitative RT-PCR analyses for OPA1 and Mfn1 and Western blot analysis for total OPA1 (L-OPA1 & S-OPA1), and Mfn1 and 2 in the retinas of AKAP1^-/-^ and WT mice. (b-d) EM and EM tomography analyses for mitochondrial fission and loss in AKAP1^-/-^ RGC somas. (b) 1.6-nm thick slices through the middle of tomographic volumes of WT RGC somas show an abundance of larger mitochondria. In contrast, the mitochondria are smaller with a greater number in AKAP1^-/-^ RGC somas, but show no alteration of cristae architecture. (c) Quantitative analyses of mitochondrial volume density, number and cross-sectional area and cristae abundance in WT and AKAP1^-/-^ RGC somas. Note that AKAP1^-/-^ RGC somas show significant loss of mitochondrial mass and cross-sectional area, and a significant increase in mitochondrial number. However, there was no significant difference in cristae abundance of AKAP1^-/-^ RGC somas. (d) Top view of the surface-rendered volume showing 3 adjacent mitochondria from a WT tomographic volume (left) and an AKAP1^-/-^ volume (right). Comparing the side view with the top view provides perspective on the distribution of the predominantly tubular cristae (in an assortment of colors) and reveals substantial heterogeneity of cristae size. The mitochondrial outer membranes were made translucent to better visualize the cristae. The number of cristae in each mitochondrion is indicated. Mean ± SD (a) and Mean ± SEM (c); *n* = 3 (a), *n* = 11 (mitochondrial volume, number and cristae abundance for WT and AKAP1^-/-^, c) and *n* = 22 or 39 (mitochondrial size for WT or AKAP1^-/-^, c); *\*P* < 0.05 and \**\*P* < 0.01 (two-tailed unpaired Student’s *t*-test). Scale bars: 200 nm.

More specifically, small isoform (80 kDa) of OPA1 protein expression was significantly decreased, whereas there was no significant difference in the large isoform (100 kDa) of OPA1 protein expression in AKAP1^-/-^ retina (Fig. 5a). We also observed a significant decrease of *Mfn1* gene expression but a significant increase of Mfn1 protein expression, whereas there was no significant difference in Mfn2 protein expression in AKAP1^-/-^ retina (Fig. 5a). Using EM and electron microscope (EM) tomography analyses, we further assessed changes of mitochondrial ultrastructure by measuring mitochondrial volume density, number, cross-sectional area and cristae abundance in AKAP1^-/-^ RGC somas. In comparison with WT mice, representative volumes generated by EM tomography provided evidence for mitochondrial fission and loss of mitochondrial mass (Fig. 5, b-d). Quantitative analysis showed significant decreases of mitochondrial volume density (13 ± 0.7%), and mitochondrial cross-sectional area (0.14 ± 0.02 µm^2^), but a significant increase of mitochondrial number (0.92 ± 0.06 per µm^2^) in AKAP1^-/-^ RGC somas compared with WT (22 ± 2 %, mitochondrial volume density; 0.20 ± 0.01 µm^2^, mitochondrial cross-sectional area; 0.67 ± 0.05 per µm^2^, mitochondrial number) (Fig. 5c). However, there was no significant difference in mitochondrial cristae abundance, which is the ratio of cristae membrane surface area divided by the mitochondrial outer membrane surface area, between WT and AKAP1^-/-^ RGC somas (Fig. 5c). Using three-dimensional (3D) reconstruction of tomographic volume, we observed that AKAP1^-/-^ mitochondria are significantly smaller than the WT mitochondria. Smaller mitochondria should have fewer cristae, given that cristae shape doesn’t change a lot, but the density of cristae was about the same between WT and AKAP1^-/-^ (Fig. 5d). These results suggest that AKAP1 loss induces an imbalance of mitochondrial dynamics by triggering mitochondrial fission and loss in RGCs.

AKAP1 loss promotes mitochondrial abnormalities and mitophagy in cardiac injury^46^. Based on our current findings of mitochondrial fission and loss, we determined whether AKAP1 loss induces mitophagosome formation. Indeed, AKAP1 loss significantly enhanced LC3-II protein expression but decreased p62 protein expression in AKAP1^-/-^ retina (Fig. 6a). Our results showed strong LC3 immunoreactivities in RGC somas and axons in the GCL of AKAP1^-/-^ mice (Fig. 6b). Notably, segmented volumes from 3D tomographic reconstructions using EM tomography revealed examples of degraded mitochondria with severe cristae depletion that were engulfed in mitophagosomes in RGC soma of AKAP1^-/-^ mice (Fig. 6c-j).

**Fig. 6.**
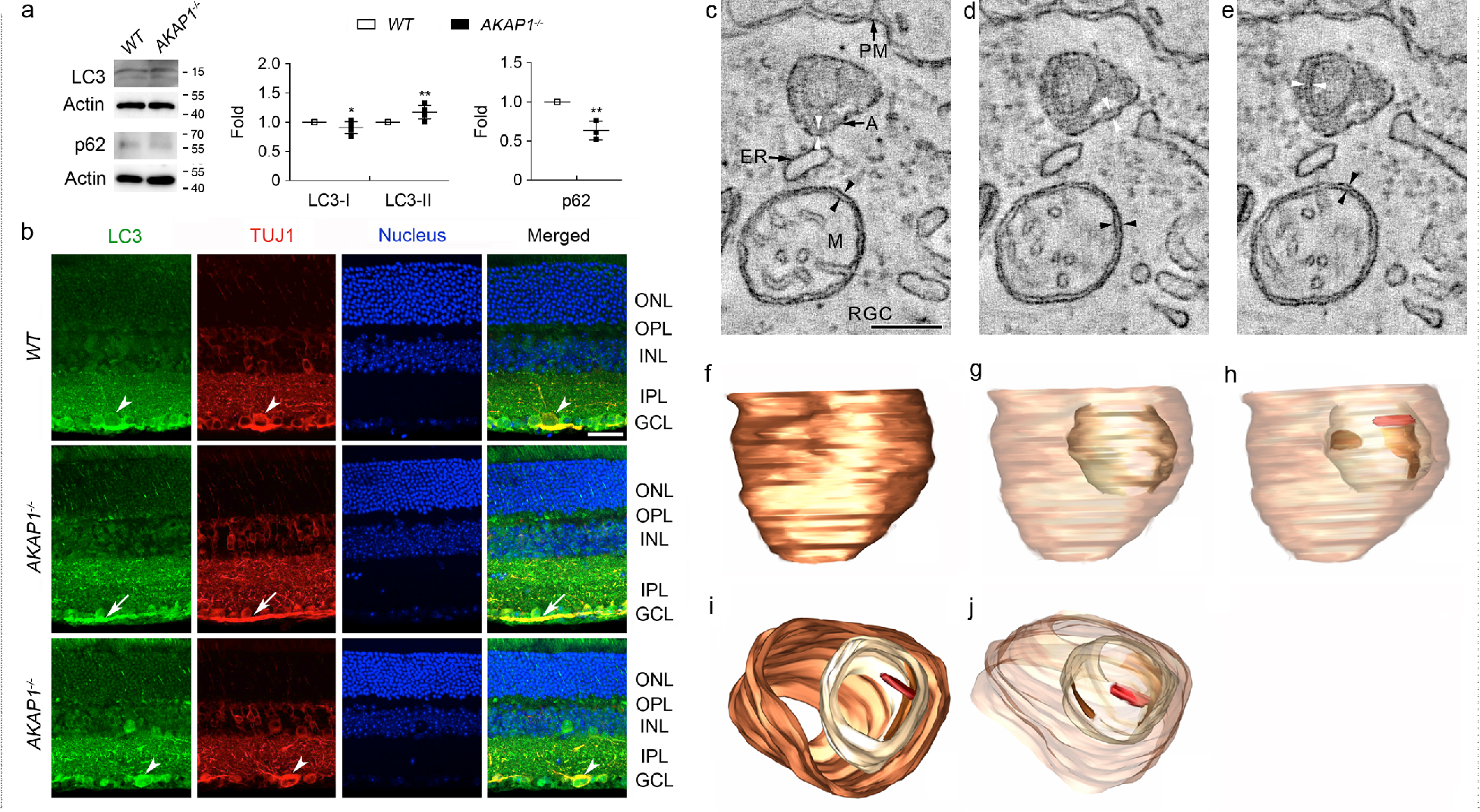
Mitophagosome formation in AKAP1^-/-^ RGC somas. (a) Western blot analyses for LC3 and p62 in the retinas of AKAP1^-/-^ and WT mice. (b) Representative images from immunohistochemical analyses for LC3 (green) co-labeled with TUJ1 (red) in RGCs and their axons. Note that LC3 immunoreactivity was localized in RGC somas (arrowheads) and their axons (arrows) in WT and AKAP1^-/-^ retina. Blue color indicates nucleus. GCL, ganglion cell layer; IPL, inner plexiform layer; INL, inner nuclear layer; OPL, outer plexiform layer; ONL, outer nuclear layer. Scale bar, 20 µm. (c-j) EM tomography and 3D reconstruction analysis. (c) A 1.6 nm-thick slice through the tomographic volume of the RGC showing an autophagosome with a degenerating mitochondrial fragment inside. A mitochondrion is nearby with a strand of ER positioned in between the autophagosome and mitochondrion. ER is typically involved with mitochondrial fission. The double membrane of the autophagosome is only partially visualized and so is indicated by white arrowheads on opposite sides. For comparison, the double membrane of the mitochondrial periphery is indicated by black arrowheads on opposite sides to emphasize that the spacing between individual autophagosomal membranes is about the same as the spacing between individual mitochondrial peripheral membranes, as it should be. Labels: A, autophagosome; M, mitochondrion; PM, RGC plasma membrane; ER, endoplasmic reticulum strand. Scale = 200 nm. (d) Another slice through the RGC volume showing a small stretch of clearly separated components of the autophagosomal double membrane (white arrowheads). For comparison, the spacing of the mitochondrial double membrane is indicated by the black arrowheads. (e) A third slice through the RGC volume that best shows the double membrane of the mitochondrial fragment (white arrowheads) inside the autophagosome. Perhaps the reason that less than half the mitochondrial double membrane is seen is because it was being degraded at the time the tissue was fixed. The spacing between the individual membranes of the fragment is the same as the spacing between the individual membranes of the mitochondrion (black arrowheads). (f) Side view with the mitophagosome membrane made transparent to see the engulfed mitochondrion. (g) Side view with the outer mitochondrial membrane made transparent to see the 2 fragments of the inner boundary membrane (IBM) and the one crista that remain. (h) Top view showing the crista and right-hand portion of the IBM inside the mitochondrion, which is inside the mitophagosome. (i and j) An oblique view with the mitophagosome and outer mitochondrial membranes made transparent to see the crista and two IBM fragments. Note that mitochondria in the AKAP1^-/-^ RGC soma demonstrate autophagosome/mitophagosome formation. Mean ± SD; *n* = 3 or 4 (a); *\*P* < 0.05 and *\*\*P* < 0.01 (two-tailed unpaired Student’s *t*-test). Scale bar: 500 nm

### AKAP1 loss results in OXPHOS dysfunction and induces oxidative stress in the retina

Since AKAP1 deletion increased superoxide production in primary hippocampal and cortical neurons in response to glutamate excitotoxicity as well as resulted in OXPHOS complex (Cx) II dysfunction^24^, we determined whether loss of AKAP1 alters OXPHOS Cxs and induces oxidative stress in the retina. We further observed that AKAP1 loss significantly increases OXPHOS Cx II but decreased OXPHOS Cx III-V protein expression in AKAP1^-/-^ retina (Fig. 7a). However, there was no significant difference in OXPHOS Cx I protein expression between WT and AKAP1^-/-^ retinas (Fig. 7a). We observed a significant increase of superoxide dismutase 2 (SOD2) protein expression in AKAP1^-/-^ retina (Fig. 7b). We also observed that SOD2 protein expression was increased in the inner retina, including RGCs in the GCL, of AKAP1^-/-^ mice (Fig. 7c and d). These results suggest that AKAP1 loss compromises OXPHOS and may contribute to oxidative stress.

**Fig. 7.**
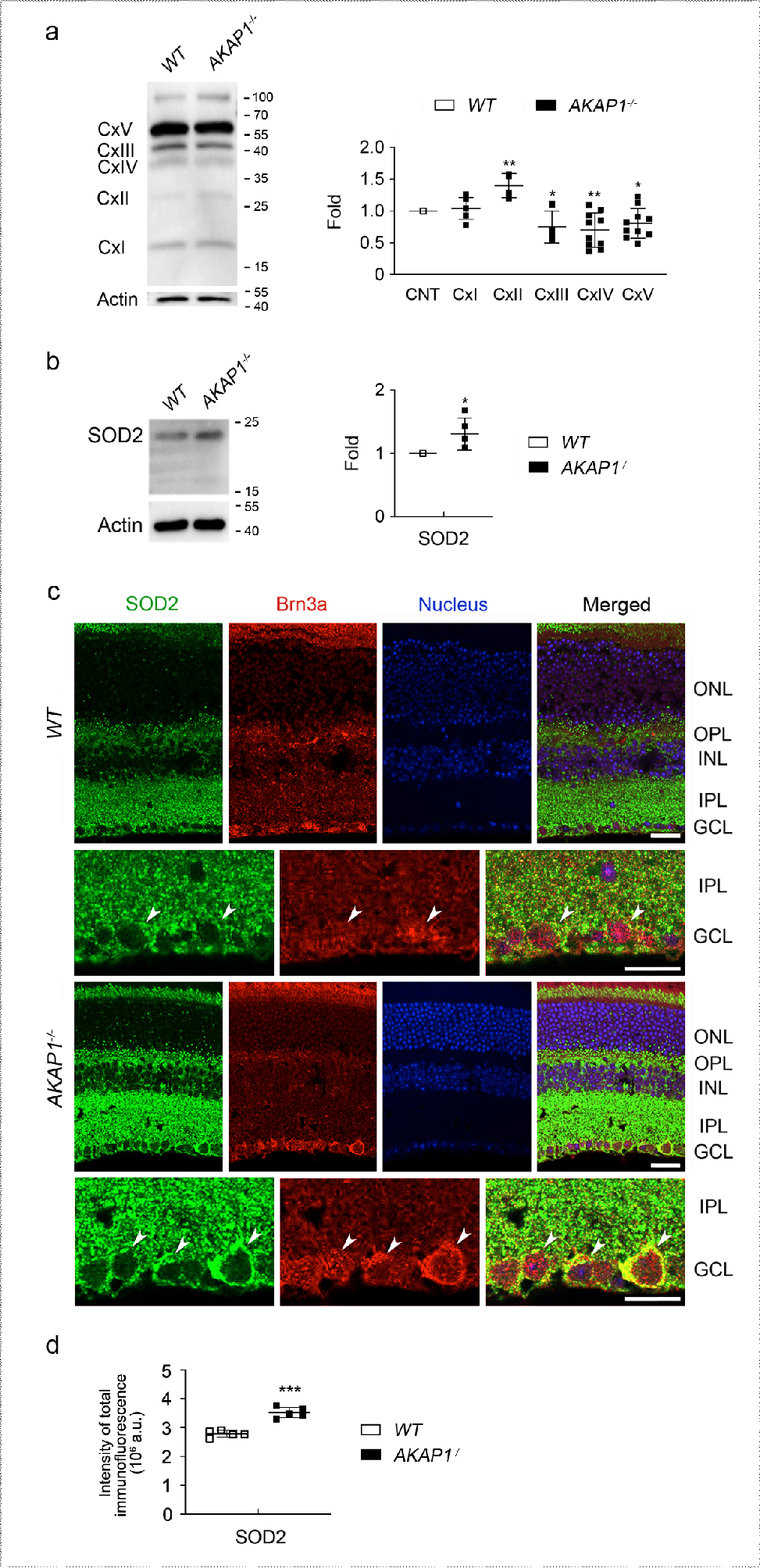
OXPHOS dysfunction and oxidative stress in AKAP1^-/-^ retinas. (a) Western blot analyses for OXPHOS Cxs in the retinas of AKAP1^-/-^ and WT mice. (b) Western blot analyses for SOD2 in the retinas of AKAP1^-/-^ and WT mice. (c) Representative images from immunohistochemical analyses for SOD2 (green, arrowheads) co-labeled with Brn3a (red, arrowheads) in the retina. Note that AKAP1^-/-^ retina shows increases of SOD2 protein expression in the inner retinal layer compared with WT retina. Blue color indicates nucleus. (d) Quantitative analysis for fluorescent intensity showed a significant increase of SOD2 immunoreactivity in the retina of AKAP1^-/-^ mice. GCL, ganglion cell layer; IPL, inner plexiform layer; INL, inner nuclear layer; OPL, outer plexiform layer; ONL, outer nuclear layer. Mean ± SD; *n* = 4 or 10 (a), *n* = 4 (b) and *n* = 10 (d). *\*P* < 0.05, *\*\*P* < 0.01 and *\*\*\*P* < 0.001 (two-tailed unpaired Student’s *t*-test). Scale bar: 20 µm.

### AKAP1 loss inactivates Akt and activates Bim/Bax pathway in the retina

We examined the expression levels of Akt protein expression, and Akt phosphorylation at serine 473 (Ser473) and threonine 308 (Thr308) in AKAP1^-/-^ retina. We observed a significant increase of total Akt protein expression but a significant dephosphorylation Akt at Ser473 and Thr308 protein expression in AKAP1^-/-^ retina (Fig. 8a). Consistently, our results showed that Akt immunoreactivity was increased in the INL and GCL of AKAP1^-/-^ retina (Fig. 8b and c). More specifically, increased Akt immunoreactivity was prominent in Brn3a-positive RGCs of AKAP1^-/-^ mice (Fig. 8b and c). Because Akt inactivation is associated with Bim/Bax activation and intrinsic apoptotic cell death^47, 48^, we further determined whether AKAP1 loss activates Bim/Bax in the retina. We observed significant increases of Bim and Bax protein expression in AKAP1^-/-^ retina (Fig. 8d). In addition, we observed that Bax immunoreactivity was increased in the GCL of AKAP1^-/-^ retina (Fig. 8d). More specifically, increased Bax immunoreactivity was prominent in RBPMS-positive RGCs of AKAP1^-/-^ mice (Fig. 8e and f).

**Fig. 8.**
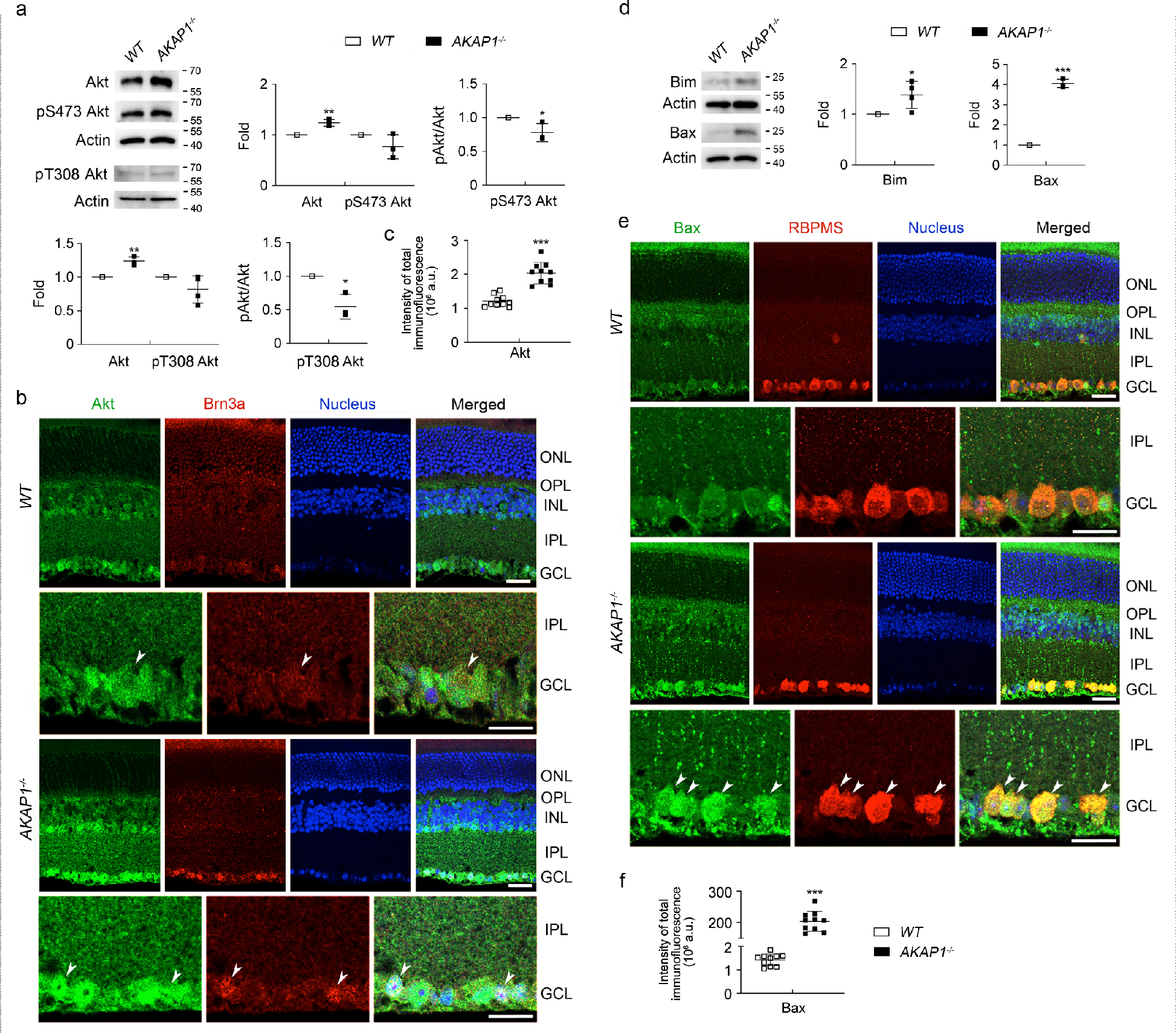
Akt inactivation and Bim/Bax activation in AKAP1^-/-^ retinas. (a) Western blot analyses for Akt, phospho-Akt Ser473 and phospho-Akt Thr308 in the retinas of AKAP1^-/-^ and WT mice. (b) Representative images from immunohistochemical analyses for Akt (green, arrowheads) co-labeled with Brn3a (red, arrowheads) in the retina. Note that AKAP1^-/-^ retina show increases of Akt protein expression in the INL and GCL compared with WT retina. Blue color indicates nucleus. (c) Quantitative analysis for fluorescent intensity showed a significant increase of Akt immunoreactivity in the retina of AKAP1^-/-^ mice. (d) Western blot analyses for Bim and Bax in the retinas of AKAP1^-/-^ and WT mice. (e) Representative images from immunohistochemical analyses for Bax (green, arrowheads) co-labeled with RBPMS (red, arrowheads) in the retina. Note that AKAP1^-/-^ retina show increases of Bax protein expression in the GCL compared with WT retina. Blue color indicates nucleus. (f) Quantitative analysis for fluorescent intensity showed a significant increase of Bax immunoreactivity in the retina of AKAP1^-/-^ mice. GCL, ganglion cell layer; IPL, inner plexiform layer; INL, inner nuclear layer; OPL, outer plexiform layer; ONL, outer nuclear layer. Mean ± SD; *n* = 3 or 4 (a and d) and *n* = 10 (c and f). *\*P* < 0.05, *\*\*P* < 0.01 and *\*\*\*P* < 0.001 (two-tailed unpaired Student’s *t*-test). Scale bar: 20 µm.

## Discussion

We here demonstrated for the first time that RGCs showed AKAP1 immunoreactivity in D2-*Gpnmb+* mice and elevated IOP triggered loss of AKAP1 in glaucomatous RGCs. In addition, elevated IOP increased CaN and total Drp1 levels in glaucomatous RGCs. It also impaired Drp1 phosphorylation at Ser637 in glaucomatous retina. These results suggest that AKAP1 and Drp1 phosphorylation at Ser637 have critical roles in RGC mitochondria and survival against glaucomatous damage. We also observed that loss of AKAP1 increased CaN and total Drp1 levels in RGCs, and compromised Drp1 phosphorylation at Ser637 in the retina, leading to mitochondrial fission and loss, and mitophagosome formation in RGCs. In addition, loss of AKAP1 resulted in OXPHOS dysfunction, as well as Akt inactivation and Bim/Bax activation; these effects contribute to glaucomatous neurodegeneration.

Impairment of mitochondrial dynamics is critically involved in glaucomatous RGCs and their axon degeneration^8,9,11, 36^, and inhibition of Drp1 rescues RGCs and their axons by preserving mitochondrial integrity in experimental glaucoma^11, 36^. Our previous study demonstrated that elevated IOP showed a significant increase of total Drp1 level, while it did not alter phosphorylation of Drp1 at Ser616 in glaucomatous retina^11^; these effects suggest that elevated IOP-mediated mitochondrial fission activity is not dependent on Drp1 phosphorylation at Ser616 in glaucomatous RGC degeneration. Hence, our novel finding that glaucomatous RGCs lacking AKAP1 trigger CaN-mediated dephosphorylation of Drp1 at Ser637 suggest an important rationale to further investigate the impact of AKAP1 loss, mitochondrial fission and its related signaling pathway in RGCs.

AKAP1 is an important regulator of mitochondrial function by PKA anchoring and Drp1 phosphorylation at Ser637, and overexpression of AKAP1 is neuroprotective in neuronal cells against the mitochondrial toxin rotenone^20, 21, 23, 24, 43^. Emerging evidence showed that mice lacking AKAP1 exhibit increased infarct following transient cerebral ischemia as well as increased Drp1 localization to mitochondria from the forebrain tissues and decreased Drp1 phosphorylation at Ser637^24^. Our findings of enhanced CaN-mediated Drp1 phosphorylation at Ser637, and extensive mitochondrial fission and loss in the retina of AKAP1^-/-^ mice suggest that AKAP1 loss directly contributes to Drp1 phosphorylation-dependent mitochondrial fission and loss in glaucomatous RGCs. This notion is also supported by the evidence of increasing LC3-II and decreasing p62 levels, as well as mitophagosome formation that contains degrading mitochondria in RGC somas of AKAP1^-/-^ mice. In addition, it is supported by the evidence of our previous report showing autophagosome/mitophagosome formation in RGC axons of glaucomatous DBA/2J mice^11^.

AMPK is a nutrient and energy sensor that maintains energy homeostatsis^49^ and mediates mitochondrial fission in response to energy stress^50^. It has been reported that there was evidence of AMPK activation by increasing AMPK phosphorylation at Thr172 in the retina and optic nerve in glaucomatous DBA/2J mice^14, 51^. Also, it has been proposed that AMPK uncouples PKA from D-AKAP1 (AKAP140/149 and other splice variants AKAP121, sAKAP84) to promote mitochondrial fission and mitophagy^52^. Furthermore, AMPK is associated with the regulation of Drp1 phosphorylation^53, 54^. In the current study, our findings showed no significant evidence of AMPK activation, suggesting that AKAP1 loss may not directly contribute to AMPK activation in the retina. Because energy stress activates AMPK^14, 55^, it is also possible that AKAP1 loss may not trigger an excessive energy stress and substantial energy deficit in the retina. In the line of this notion, our results showed that the level of dephosphorylation of Drp1 Ser637 in AKAP1^-/-^ retina was relatively modest compared to the level of dephosphorylation of Drp1 Ser637 in glaucomatous retina. Hence, our results intriguingly raise a possibility that combined induction of both AMPK activation and AKAP1 loss may exacerbate mitochondrial fission and loss in glaucomatous RGCs and their axons, leading to the promotion of energy crisis and cell death. We believe that further studies will be required to investigate the relationship between AMPK activation and AKAP1 loss in glaucomatous RGCs.

The increase in LC3-II accumulates autophagosomes, but does not guarantee autophagic degradation. If, however, the amount of LC3-II further accumulates in the presence of lysosomal protease inhibitors, this would indicate enhancement of autophagic flux^56^. On the other hand, the protein expression level of p62/SQSTM1, a polyubiquitin binding protein, serves as autophagic receptor and is a good indicator of autophagic flux^57–59^. Also, p62 protein expression has been used as autophagic marker with LC3-II^60^. Although p62 level alone is not indicative of activation of autophagy, p62 is also a marker for the degradation phase of autophagosomes and has a lot of functions beyond autophagy^59^. In the current study, our findings showed that loss of AKAP1 increased LC3-II protein expression but decreased p62 protein expression in AKAP1^-/-^ retina. Intriguingly, AKAP1^-/-^ heart reduced p62 protein expression and enhanced mitophagy and apoptosis^46^. Collectively, our results suggest that the increased LC3-II and decreased p62 contribute to the induction of autophagy/mitophagy in AKAP1^-/-^ retina. Hence, we raise a possibility that loss of AKAP1 may not only induce autophagosome/mitophagosome formation, but also trigger increase of autophagic flux and degradation in RGCs. Also, whether this autophagic/mitophagic process is a compensatory or pathological pathway in AKAP1^-/-^ retina should be considered in the future studies.

Previous study indicated the notion that AKAP1 regulates mitochondrial dynamics by posttranslational modifications, but levels of Drp1, OPA1 and Mfn2 were unaltered in total forebrain homogenates from AKAP1^-/-^ mice^24^. Of interest, however, we observed a decrease of total OPA1, but an increase of total Drp1 and Mfn1 in the retina of AKAP1^-/-^ mice. Inner membrane-anchored long OPA1 (L-OPA1) undergoes proteolytic cleavage resulting in short OPA1 (S-OPA1)^61^. Cells contain a mixture of L-OPA1, required for mitochondrial fusion and S-OPA, which facilitate mitochondrial fission under normal conditions. Disruption of this balance in times of stress, result in mitochondrial fragmentation^61^. Thus, these unexpected, but important results propose the possibility that AKAP1 may contribute to the regulation of mitochondrial dynamics not only by posttranslational modifications, but also by alteration of fusion/fission protein expression in the retina. Future studies will be needed to clarify the molecular mechanisms underlying the alteration of fusion/fission proteins in the retina lacking AKAP1.

Emerging evidence showed that AKAP1 deletion results in OXPHOS Cx II dysfunction in neuronal cells^24^. OXPHOS Cx II is a source of the mitochondrial reserve respiratory capacity that is regulated by metabolic sensor and promotes cell survival against hypoxia. In the present study, the observation that increasing OXPHOS Cx II, but decreasing OXPHOS Cxs III-V activities triggered by loss of AKAP1 in the retina is likely to imply that AKAP1 plays a critical role in OXPHOS function in RGCs. Moreover, increasing OXPHOS Cx II activity may be an important endogenous compensatory defense mechanism in response to oxidative stress.

Further, this imbalance of OXPHOS Cxs suggests another possibility that AKAP1 loss may contribute to an increase of ROS production and decrease of ATP production in the retina that are previously reported in pressure and/or oxidative stress-induced RGC degeneration^11^. In particular, association of POAG with polymorphism of mitochondrial cytochrome c oxidase subunit 1 of OXPHOS Cx IV suggests a potential role of abnormal OXPHOS-mediated mitochondrial dysfunction in glaucoma pathogenesis^3, 4^. Therefore, these results collectively suggest that AKAP1 loss in glaucomatous retina contributes to OXPHOS dysfunction that is associated with metabolic and oxidative stress by increasing ROS production and decreasing ATP reduction, leading to RGC death during glaucomatous neurodegeneration.

Loss of AKAP1 inactivates Akt signaling and increases apoptosis in cardiac dysfunction^62, 63^. Akt pathway controls the expression of apoptosis-related gene through transcription factors such as FoxO (FoxO1 and FoxO3a)^64, 65^ and Akt activation promotes cell survival by inhibiting FoxO3a/Bim/Bax pathway^48, 64, 65^. The pro-apoptotic Bcl-2 homology domain 3-only protein Bim induces apoptosis, primarily through its increased binding activity toward multiple pro-survival Bcl-2-like proteins, whose dissociations activate Bak and Bax^66, 67^. In addition, Bax-mediated mitochondrial outer membrane permeabilization is an important pathophysiological mechanism for metabolic dysfunction and cell death^68, 69^, and Bax activation plays a critical role in mitochondrial dysfunction-mediated RGC degeneration^11, 12^. Because Drp1 contributes to Bax oligomerization, mitochondrial fission and the cellular apoptotic pathway^70–72^, increased total Drp1 and Bax levels in AKAP1^-/-^ retinas suggest that AKAP1 loss may contribute to Drp1-mediated Bax oligomerization, leading to mitochondrial fission. Recent evidence from our group and others indicated that activation of cAMP/PKA pathway promotes RGC survival ^48, 73–75^ and that AKAP1-mediated neuroprotection requires PKA anchoring and Drp1 phosphorylation at Ser637^21, 23, 24, 43, 76^. In this regard, our observation that AKAP1^-/-^ retinas dephosphorylates Akt at Ser473 and SerT308, and activates Bim/Bax pathway raise the possibility that AKAP1 may be critical to RGC protection by modulating Akt/Bim/Bax pathway against glaucomatous damage. Future studies will be necessary to examine whether overexpression of AKAP1 rescues RGCs by promoting Drp1 phosphorylation at S637, maintaining metabolic activity, activating Akt pathway and inhibiting Bim/Bax pathway.

In summary, our results represent the first direct evidence of a Drp1 phosphorylation-mediated mitochondrial pathogenic mechanism that leads to mitochondrial fission and metabolic dysfunction in glaucomatous RGC degeneration. Also, we provide evidence for the first time that AKAP1 loss-mediated CaN activation and Drp1 dephosphorylation at Ser637 result in mitochondrial fission, mitophagosome formation, OXPHOS dysfunction and oxidative stress, as well as Akt inactivation and Bim/Bax activation in RGCs. Therefore, we propose overexpression of AKAP1 and modulation of Drp1 phosphorylation at Ser637 as therapeutic strategies for neuroprotective intervention in glaucoma and other mitochondria-related optic neuropathies.

## Materials and methods Animals

Adult female DBA/2J and D2-*Gpnmb^+^* mice (The Jackson Laboratory), and adult male WT and AKPA1^-/-^ mice were housed in covered cages, fed with a standard rodent diet ad libitum, and kept on a12 h light/12 h dark cycle. Animals were assigned randomly to experimental and control groups. The following PCR primers were used to determine AKAP1 genotype: GGAGGCGATCACAGCAACAACCG (R), ATACAGAAGCAGATCACTCAGGAGG (F-WT), CAGTCCCAAGGCTCATTTCAGGCC (F-KO)^24^. To investigate the effect of IOP elevation, 20 10 month-old DBA/2J and 10 age-matched D2-*Gpnmb^+^* mice were used. To investigate the effect of AKAP1 loss, 10 10 month-old AKAP1^-/-^ and 10 age-matched WT mice were used. All procedures concerning animals were in accordance with the Association for Research in Vision and Ophthalmology Statement for the Use of Animals in Ophthalmic Vision Research and under protocols approved by Institutional Animal Care and Use Committee at the University of California, San Diego.

### Tissue preparation

Mice were anesthetized by an intraperitoneal (IP) injection of a cocktail of ketamine (100 mg/kg, Ketaset; Fort Dodge Animal Health) and xylazine (9 mg/kg, TranquiVed; VEDCO Inc.) prior to cervical dislocation. The investigators were not blinded during tissue and data collection from mice. Blinding was used during data analysis. For immunohistochemistry, the retinas were dissected from the choroids and fixed with 4% paraformaldehyde (Sigma-Aldrich) in phosphate buffered saline (PBS, pH 7.4) for 2 h at 4 °C. Retinas were washed several times with PBS then dehydrated through graded levels of ethanol and embedded in polyester wax. For Western blot and PCR analyses, extracted retinas were immediately used.

### Western blot analyses

Harvested retinas (*n* = 4 retinas/group) were homogenized for 1 min on ice with a modified RIPA lysis buffer (150 mM NaCl, 1 mM EDTA, 1% NP-40, 0.1% SDS, 1 mM DTT, 0.5% sodium deoxycholate and 50 mM Tris-Cl, pH 7.6), containing complete protease inhibitors. The lysates were then centrifuged at 15,000 g for 15 min and protein amounts in the supernatants were measured by Bradford assay. Proteins (10-20 µg) were separated by SDS/PAGE and electrotransferred to polyvinylidene difluoride membranes (GE Healthcare Bio-Science). The membranes were blocked with 5% non-fat dry milk and PBS/0.1% Tween-20 (PBS-T) for 1 h at room temperature and incubated with primary antibodies for overnight at 4 °C. Primary antibodies are mouse monoclonal anti-Actin antibody (1:20,000; Millipore; Cat#, MAB1501), rabbit monoclonal anti-AKAP1 (1:1000; Cell Signaling; Cat# 5203), rabbit polyclonal anti-Akt antibody (1:5000; Cell Signaling; Cat# 92725), mouse monoclonal anti-phospho-Akt Ser473 antibody (1:100, Cell Signaling; Cat# 9271), rabbit monoclonal anti-phospho-Akt Thr308 antibody (1:1000, Cell Signaling; Cat# 4056), rabbit polyclonal anti-AMPK*α*1/2 antibody (1:1000, Santa Cruz Biotechnology; Cat# sc25792), rabbit monoclonal anti-AMPK*α* Thr172 antibody (1:1000, Cell Signaling Cat# 2535), mouse monoclonal anti-Bax antibody (6A7; 1:500; Santa Cruz Biotechnology; Cat# sc-23959), rabbit polyclonal anti-Bim antibody (1:100, Santa Cruz Biotechnology; Cat# sc-8265), mouse monoclonal anti-CaN antibody (1:200, BD Bioscience; Cat# 610259), mouse monoclonal anti-Drp1 antibody (1:1000; BD Biosciences; Cat# 611113), rabbit polyclonal phospho-Drp1 Ser637 antibody (1:750; Cell Signaling; Cat# 4867), rabbit polyclonal phospho-Drp1 Ser616 antibody (1:750; Cell Signaling; Cat# 3455), rabbit polyclonal anti-LC3 antibody (1:2000; MBL International; Cat# PM036), mouse monoclonal anti-Mfn1 antibody (1:20000; Abcam; Cat# ab57602), mouse monoclonal anti-Mfn2 antibody (1:2000; Abcam; Cat# ab56889), mouse monoclonal anti-OPA1 antibody (1:2000; BD Biosciences; Cat# 612607), rat/mouse monoclonal anti-OXPHOS Cx Cocktail Kit (1:1000; Invitrogen; Cat# 458099), rabbit polyclonal p62 antibody (1:1000; MBL International; Cat# PM045) and rabbit polyclonal anti-SOD2 antibody (1:5000; Santa Cruz; Cat# sc-30080).

Membrane were washed several times with PBS-T then incubated with peroxidase-conjugated goat anti-mouse or rabbit IgG (1:1000-7000; Bio-Rad; Cat# 1721011 or 1706515) for 1 h at room temperature and developed using enhanced chemiluminescence substrate system. The images were captured and quantified by using ImageQuant™ LAS 4000 system (GE Healthcare Bio-Science) and band densities were normalized to that of actin, total Drp1, total Akt, or total AMPK.

### Immunohistochemistry

Immunohistochemical staining of 7 μm wax sections of full thickness retina were performed. Sections from wax blocks from each group (*n* = 4 retinas/group) were used for immunohistochemical analysis. Primary antibodies included rabbit monoclonal anti-AKAP1 antibody (1:100, Cell Signaling; Cat# 5203), rabbit polyclonal anti-Akt antibody (1:100, Cell Signaling; Cat# 92725), mouse monoclonal anti-Bax antibody (6A7; 1:50, Santa Cruz Biotechnology; Cat# sc-23959), goat polyclonal anti-Brn3a antibody (1:500, Santa Cruz Biotechnology; Cat# sc-23959), mouse monoclonal anti-CaN (1:200, BD Bioscience; Cat# 610259), mouse monoclonal anti-Drp1 antibody (1:25; BD Biosciences; Cat# 611113), rabbit polyclonal anti-Drp1 (1:50, Santa Cruz Biotechnology, Cat# sc-32898), rabbit polyclonal anti-LC3 antibody (1:2000; MBL International); Cat# PM036, rabbit polyclonal anti-RBPMS (1:300, Novus; Cat# NBP2-20112), rabbit polyclonal anti-SOD2 antibody (1:300, Santa Cruz Biotechnology; Cat# sc-30080) and mouse monoclonal anti-TUJ1 (1:300, BioLegend; Ct# 801202). To prevent non-specific background, tissues were incubated in 1% bovine serum albumin/PBS for 1 h at room temperature before incubation with the primary antibodies for 16 h at 4°C. After several wash steps, the tissues were incubated with the secondary antibodies, Alexa Fluor 488 or 568 dye-conjugated donkey anti-mouse IgG (1:100, Invitrogen; Cat# R37114 or A10037), Alexa Fluor 488 or 568 dye-conjugated donkey anti-rabbit IgG (1:100, Invitrogen; Cat# R37118 or A10042) and Alexa Fluor 568 dye-conjugated donkey anti-goat IgG (1:100, Invitrogen; Cat# A11057) for 4 h at 4°C and subsequently washed with PBS. The sections were counterstained with the nucleic acid stain Hoechst 33342 (1 μg/mL; Invitrogen) in PBS. Images were acquired with confocal microscopy (Olympus FluoView1000; Olympus, Tokyo, Japan). Each target protein fluorescent integrated intensity in pixel per area was measured using the ImageJ software^77–79^. All imaging parameters was remained the same and corrected with the background subtraction.

### RT-PCR and quantitative real-time RT-PCR

Total RNA was isolated from retinas of each group (*n* = 4 retinas/group) using the Absolutely RNA Miniprep Kit (Stratagene), according to the manufacturer’s protocol as previously described^48^. cDNAs were synthesized from total RNAs with iScript cDNA Synthesis Kit (Bio-Rad) according to the manufacturer’s protocols. Reverse Transcription (RT)-PCR was performed with cDNAs synthesized from 1 µg of the total RNA of each group as a template and specific primers for: AKAP1, CTGCCAGTCAGTACTCAGCC (F), CTTTGGCACCTCGATCTCCC (R), OPA1, CATCTACCTTCCAGCTGCCC (F), TCTCCTCCTTCACAGCCTCC (R), and Mfn1, GTTTTCCCTGGGCTGGTCTT (F), TTTTCCAAATCACGCCCCCA (R). For the quantification of the relative mRNA expressions of each group, real-time PCR was carried out using MX3000P real-time PCR system (Stratagene) as follows. cDNAs were amplified using SSO Adv. Universal SYBR Green SuperMix (Bio-Rad) and the specific primers for 40 cycles (95 °C for 5 min, and 40 cycles (95 °C for 30 sec, 60 °C for 30 sec, and 72 °C for 30 sec)). Output data were obtained as *Ct* values and the differential mRNA expression of the target genes among groups was calculated using the comparative *Ct* method. GAPDH mRNA, an internal control, was amplified along with the target gene, and the *Ct* value of GAPDH was used to normalize the expression of the target gene.

### Transmission electron microscopy

For conventional EM, two eyes from each group (*n* = 2 mice) were fixed via cardiac perfusion with 2% paraformaldehyde, 2.5% glutaraldehyde (Ted Pella, Redding, CA) in 0.15M sodium cacodylate (pH 7.4, Sigma) solution at 37°C and placed in pre-cooled fixative of the same composition on ice for 1 h. The following procedure was used to optimize mitochondrial structural preservation and membrane contrast. The retinas were dissected in a solution of 0.15M sodium cacodylate plus 3mM calcium chloride (pH 7.4) on ice and then post-fixed with a 1% osmium tetroxide, 0.8% potassium ferrocyanide, 3mM calcium chloride in 0.1M sodium cacodylate solution (pH 7.4) for 1 h, washed with ice-cold distilled water, poststained with 2% uranyl acetate solution at 4°C, dehydrated using graded ethanols, and embedded in Durcupan resin (Fluka). A similar procedure was used for cultured primary RGCs from each group.

Ultrathin (70 nm) sections were post-stained with uranyl acetate and lead salt solutions and evaluated using a JEOL 1200FX transmission EM operated at 80kV. Images were recorded on film at 8,000X magnification. The negatives were digitized at 1800 dpi using a Nikon Cool scan system, giving an image size of 4033 x 6010 pixel array and a pixel resolution of 1.77 nm. For quantitative analysis, the number of mitochondria was normalized to the total area occupied by axons in each image, which was measured using ImageJ (http://rsb.info.nih.gov/ij/).

Mitochondrial lengths were measured with ImageJ. The mitochondrial volume density, defined as the volume occupied by mitochondria divided by the volume occupied by the axoplasm, was estimated using stereology as follows. A 112 x 112 square grid (112 x 112 chosen for ease of use with Photoshop) was overlaid on each image loaded in Photoshop (Adobe Systems Inc.), and mitochondria and axoplasm lying under intercepts were counted. The relative volume of mitochondria was expressed as the ratio of intercepts coinciding with this organelle relative to the intercepts coinciding with axoplasm.

### Electron microscope tomography

Sections of retina tissues from each group were cut at a thickness of 400 nm. For each reconstruction, a double-tilt series of images at one degree tilt increment was collected with an FEI titan hibase electron microscope operated at 300 kV. Images were recorded with a Gatan 4Kx4K CCD camera. The magnification was 11,000x and the pixel resolution was 0.81 nm. The IMOD package was used for alignment, reconstruction and volume segmentation. Volume segmentation was performed by manual tracing of membranes in the planes of highest resolution with the Drawing Tools and Interpolator plug-ins^80^. The mitochondrial reconstructions and surface-rendered volumes were visualized using 3DIMOD. Measurements of mitochondrial outer, inner boundary, and cristae membrane surface areas and volumes were made within segmented volumes using IMODinfo. These were used to determine the cristae abundance, defined as the ratio: sum of the cristae membrane surface areas divided by the mitochondrial outer membrane surface area. Movies of the tomographic volume were made using IMODmovie.

## Statistical analyses

Data were shown as mean ± S.D. Comparison of experimental conditions was evaluated using the two-tailed unpaired Student’s t-test between groups. *P* < 0.05 was considered to be statistically significant.

## Acknowledgements

This work was supported, in part, by NIH grants EY018658, P30 EY022589, T32 EY026590, 5P41GM103412, and an unrestricted grant from Research to Prevent Blindness (New York, NY) and UCSD Academic Senate.

